# The City Nature Challenge – A global citizen science phenomenon contributing to biodiversity knowledge and informing local government practices

**DOI:** 10.1101/2022.11.14.516526

**Authors:** Estibaliz Palma, Luis Mata, Kylie Cohen, Doug Evans, Bernard Gandy, Nadine Gaskell, Hiliary Hatchman, Anna Mezzetti, Deborah Neumann, Jessica O’Keefe, Amy Shaw, Millie Wells, Laurence Williams, Amy K. Hahs

## Abstract

The bioblitz phenomenon has recently branched into cities, presenting exciting opportunities for local governments to channel participants’ efforts toward local issues. The City Nature Challenge (CNC) is one such initiative that has been quickly uptaken by hundreds of municipalities worldwide. Despite high participation, we still lack a framework for evaluating how the CNC contributes to local biodiversity knowledge and to inform local government practices. Here, we develop such a tool and present a case study that illustrates its applicability. We demonstrate that the collected records contributed to a better understanding of contemporary, local biodiversity patterns and provided a more realistic representation of understudied groups such as insects and fungi. Importantly, we show that the CNC presented local governments with a cost-effective tool to make informed, evidence-based management and policy decisions, improve education and engagement programs, foster cross-council collaborations, and support a stronger sense of environmental stewardship within the local community.

## Introduction

During the last two decades, the use of the term citizen science has exponentially grown within and beyond the scientific literature. This growth reflects the increasing number and diversity of initiatives that synergistically bring together professional researchers and the general public to advance knowledge across a wide array of scientific disciplines (Bonney et al. 2014, Pocock et al. 2017). Citizen science, also referred to as community science (Heigl et al. 2019), has gained worldwide traction due to an increasing interest from the general public to contribute to the scientific endeavour. Citizen scientists’ contributions have been further facilitated by calls for open science (Cribb and Sari 2010, Murray-Rust 2008) and innovations in web-based and app technologies (Newman et al. 2012). The nature and approach of citizen science projects can be very diverse, with citizen scientists being involved in some or most aspects of the project (Shirk et al. 2012). Yet, they all aim to achieve similar overall objectives. First, they aim to advance fundamental knowledge and contribute applied tools to solve pressing problems. Second, they aim to bring together input from a diversity of stakeholders in the project design and execution (Kurle et al. 2022). By doing so, they aim to facilitate a relationship among the several parts involved in managing an issue (Roger and Klistorner 2016). Research that promotes diversity of participating actors not only creates community connections, it also facilitates knowledge transfer and crowdsourcing (Roger and Klistorner 2016), and strengthens the learning experience (Hartman et al. 2019, Newman et al. 2012). Ultimately, it has the potential to catapult changes in management, policy and governance (Couvet and Prevot 2015, Lowman et al. 2019). While citizen science projects have made lasting and meaningful contributions across a wide range of disciplines (Pettibone et al. 2017), we draw attention here to projects that specifically focus on advancing knowledge of biodiversity, promoting education in biodiversity conservation, and increasing the public’s involvement with biodiversity-oriented practices at the local scale (Bonney et al. 2009, Dickinson et al. 2010, Dickinson et al. 2012, Pocock et al. 2018).

Biodiversity-based citizen science initiatives have gained enormous momentum, fuelled by people’s concerns with global environmental change, particularly current and predicted rates of habitat transformation and species loss. Often, citizen scientists are recruited as volunteers, either to work independently or alongside professional researchers, to boost the resourcing capacity of projects to collect biodiversity data across larger areas and longer time periods (Pocock et al. 2017). A key challenge is striking the balance amongst flexibility in data collection, data quality, and the number of volunteers – more elaborate data collection protocols are likely to achieve higher quality datasets, usually at the expense of smaller and less diverse groups of volunteers (Brown and Williams 2019). A notable example of biodiversity-oriented citizen science projects with well-developed sampling designs, trained volunteers, and long-term professional oversight are those led by the Cornell Ornithology Lab (Bonney et al. 2009), who have successfully built strong connections with their volunteers to study diverse aspects of bird biodiversity, ecology and conservation (Bhattacharjee 2005). Other initiatives with more relaxed data collection protocols may achieve broader participation; however, their power to answer specific research questions may be more limited. A quintessential example of initiatives with more relaxed data collection protocols are bioblitzes. These are short to mid duration biodiversity surveys, aimed at finding and identifying as many species as possible at a given location, during a specific timeframe (National Geographic Society n.d.). Bioblitzes are typically organised in natural areas of particular significance, often but not exclusively where biodiversity knowledge is scarce but highly valuable to guide local management. Despite not necessarily being driven by a specific research agenda, bioblitzes are well-known for their contribution to contemporary biodiversity knowledge (Ballard et al. 2017, Spear et al. 2017) and the documentation of species not previously known to Western science (Barrett 2015, Cassis and Symonds 2016, Fagan-Jeffries et al. 2019, Lambkin and Bartlett 2011, Vendetti et al. 2018). Importantly, bioblitzes contribute to increase participants’ engagement with nature and conservation, especially those with no previous expertise (Lundmark 2003, Postles and Bartlett 2018, Roger and Klistorner 2016). Recently, the bioblitz movement has percolated to urban environments – this is a largely underexplored space in biodiversity and education research and practice that we believe presents an exciting opportunity to advance knowledge and highlight the value of nature in cities.

Surveying biodiversity across urban environments is key to understanding and quantifying the effects of anthropogenic pressures on biodiversity (Aronson et al. 2014). Greenspaces within cities and towns support ample microbial, fungal, plant, and animal diversity (Aronson et al. 2018, Baldock et al. 2019, Gallo et al. 2017, MacGregor-Fors et al. 2016, Mata et al. 2021, Threlfall et al. 2017). Equally importantly, they provide many socio-cultural benefits to people who interact with them (Flies et al. 2017, Lai et al. 2019, Maller et al. 2019, Mata et al. 2020). Remnant bushland, public parks, and other types of greenspace typically visited by urban bioblitz participants embody the day-to-day opportunity to be in contact with nature for the majority of city dwellers. Not surprisingly, a wide range of urban stakeholders – from researchers, practitioners, built-environment professionals, conservationists, and policymakers to wildlife gardeners, Indigenous communities, ArtScience advocates, and friends-of-groups – are increasingly, and often synergistically, working towards promoting and demonstrating the benefits of urban greenspaces for both people and the rest of nature (Aronson et al. 2017, Cumpston 2020, Lepczyk et al. 2017, Mata et al. 2020, Mumaw and Mata 2022, Nilon et al. 2017, Parris et al. 2018, Renowden et al. 2022, Soanes et al. 2019). Urban bioblitzes provide an opportunity to simultaneously gather biodiversity records across greenspace networks (Rega-Brodsky et al. 2022) and strengthen the link between city dwellers and the governance of biodiversity and ecosystems in urban environments (McPhearson et al. 2016). By leading these initiatives, local governments and naturalist groups are key players to channel citizen science efforts towards specific local issues, and to promote the use of a centralised data collection repository across participants and projects (Kobori et al. 2016). In this study, we focus on the City Nature Challenge (CNC for short), because we recognise that it represents an outstanding example of a global biodiversity-oriented urban bioblitz event (Box 1).

While the CNC and other related projects (e.g. Great Southern Bioblitz) have been very successful in contributing biodiversity records, and have been readily taken up by hundreds of municipalities and shires across the world (Box 1), there is limited evidence on how data produced by urban bioblitzes are contributing to increased biodiversity knowledge at the scale of local governments. We argue that the CNC provides a unique opportunity to quantify how biodiversity knowledge changes at the local government scale. Specifically, we contend that data derived from the CNC can be easily analysed in the context of a given local government area or areas to assess the number and identity of species that (1) are likely locally extinct; (2) are currently present but have not been recently recorded; and (3) have not previously been recorded, whether because they have remained historically undetected or have only been recently introduced into the area (Figure 3). Here, we present empirical evidence to support these points using a case study from Melbourne, Australia, which embodies the power of citizen science to advance biodiversity knowledge and citizen engagement in urban environments. We then discuss how local governments have been taking up and translating new knowledge acquired during the CNC to inform and improve their urban nature and community engagement practices. Finally, we argue for ways in which the theoretical advances and empirical protocols we present in this study may be robustly transferred to inform the practices of other local government areas and discuss the benefits of scaling up our evaluation approach to regional and global scales.

### BOX 1

**City Nature Challenge**

The City Nature Challenge (CNC for short) began in the United States in 2016, when staff at the Natural History Museum of Los Angeles and the California Academy of Sciences conceived a friendly competition between San Francisco and Los Angeles to see which city could record the largest number of species by the largest number of participants during eight days (https://citynaturechallenge.org/). Over the following years, an increasing number of cities have joined this initiative (Figure 1a) and called on their residents to find and document an increasing amount of urban biodiversity (Figure 1b-d). In just six years, the CNC has evolved to be an internationally recognised urban bioblitz event. Every year, at the end of April, citizen scientists globally come together for four days to document the largest possible number of species in urban areas. By 2021, CNC initiatives spread to over 400 cities across more than 40 countries. During 2021, over 1,200,000 records of more than 45,000 species were collected by approximately 52,000 participants (Table S1). These figures, however, varied greatly across the participating countries (from 36 to 567,129 records contributed by a single country; Figure 2), with the United States and South Africa being the countries contributing the largest number of records (Table S2). Despite the competitive nature of the CNC, where urban nodes compete against each other to tally the highest count of local species, this initiative allows participants worldwide to collaborate and contribute to document biodiversity patterns at global level.

**Figure 1.**
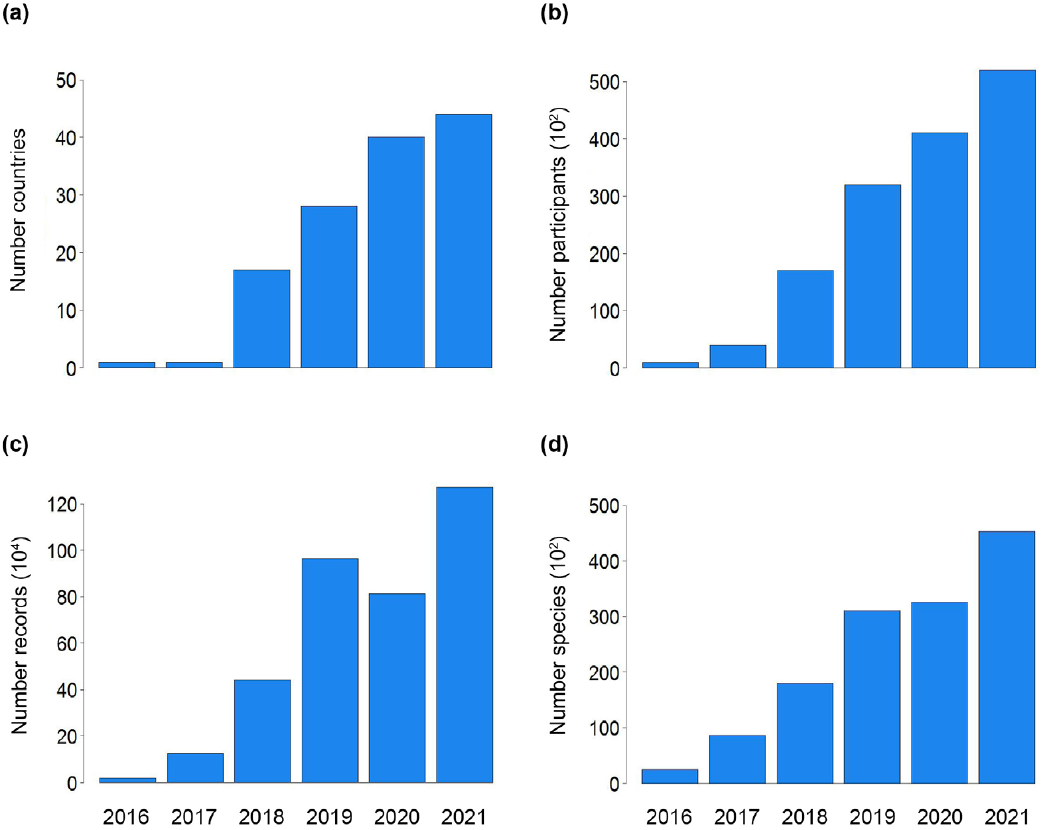
Number of countries **(a)** and participants **(b)** who took part in the City Nature Challenge from 2016 to 2021, along with records contributed **(c)** and species found **(d)** for the same time period worldwide.

**Figure 2.**
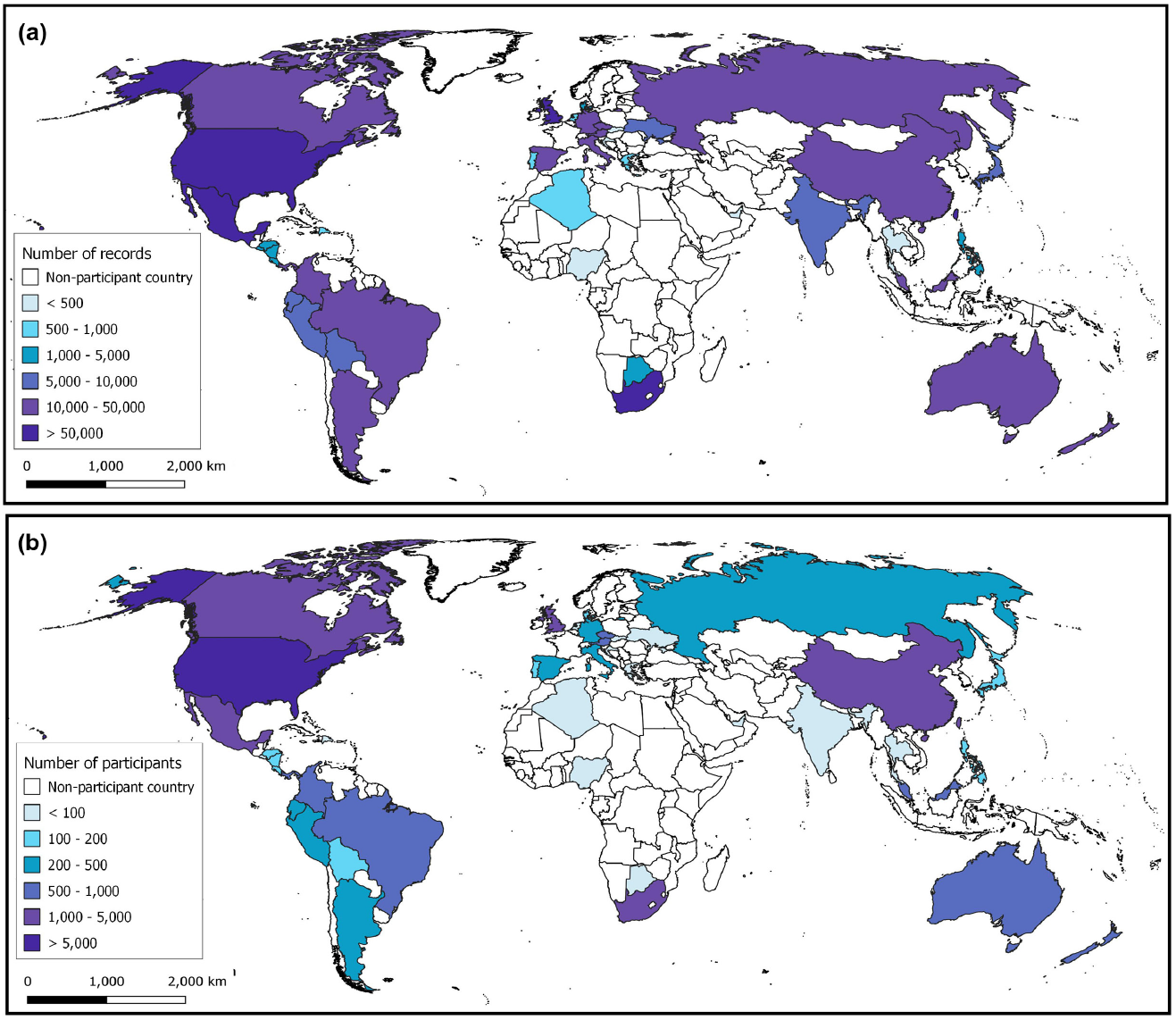
**(a)** Number of observations collected across countries that participated in the 2021 City Nature Challenge. **(b)** Number of participants by country during the 2021 City Nature Challenge.

Record collection is done through media, commonly photographs, and shared into virtual platforms – often iNaturalist (Appendix A). This is a non-destructive, open source and media-verifiable sampling approach that leads to presence-only data (i.e. absence of species is not documented directly). Once observations are uploaded to the virtual platform, they can be seen and identified by the entire virtual community within the platform. The fact that observation contribution and identification can be done simultaneously or at different times and/or by different people makes participation in the CNC open to anyone, and equally importantly, tailored to anyone’s interests and skills. It is perhaps the flexible nature of this methodology and its inclusivity regarding participation what has driven its rapid uptake worldwide.

Coordination across CNC nodes to collect observations in a common virtual platform results in large spatial coverage of biodiversity in urban areas, and provides a transparent way to compare observations across cities by anyone within or outside the CNC community. Observations collected through the CNC are openly available to download and use by anyone – scientists increasingly include urban biodiversity data collected by citizen scientists in their research (Rega-Brodsky et al. 2022). Moreover, observations that are identified to a certain degree of community consensus (e.g. *research grade* in iNaturalist) are automatically shared with other global biodiversity information platforms (e.g. Global Biodiversity Information Facility, Atlas of Living Australia). Finally, lessons learnt through CNC data are quickly absorbed by local governments and organisations to make more-informed management decisions, update policy, or create tools to deliver education programmes (see section *Lessons learnt: Uptake of City Nature Challenge findings by local government*).

### City Nature Challenge 2021: The ‘Melbourne Eastern Metropolitan Area’ node

The ‘Melbourne Eastern Metropolitan Area’ was a node of the City Nature Challenge 2021 constituted by eight Local Government Areas from eastern Melbourne, Australia (Box 2). Council staff led the organisation of the node and teamed up with local naturalist groups, learned societies, and friend groups to co-run biodiversity surveys. They also engaged academic researchers to co-design the evaluation that led to this work.

During the CNC 2021, the 291 participants of the ‘Melbourne Eastern Metropolitan Area’ node contributed 4,638 biodiversity records representing 974 taxa across nine broad taxonomic groups (Tables S3-S4). These records indicate that the participants found around 1% of the 2,112 species that had not been recorded recently – i.e. in the last three decades – for this area (Table S5, Figure 4e), therefore providing evidence these species have not gone locally extinct. Participants also documented about 10% of the 4,206 species that had been recently recorded for this area in biodiversity repositories (Table S5, Figure 4h). In addition, participants found 135 taxa that had never been recorded in this area (Figure 4g), increasing the local species richness by almost 4% (Table S5). At least 22 of these newly recorded species were introduced to Victoria (Table S3, Figure S2).

#### BOX 2

**Evaluation approach for the ‘Melbourne Eastern Metropolitan Area’ node of the 2021 City Nature Challenge**

The ‘Melbourne Eastern Metropolitan Area’ node of the 2021 CNC was born as an *ad-hoc* collaboration between the City of Boroondara, the City of Greater Dandenong, the City of Knox, the City of Manningham, the City of Maroondah, the City of Monash, the City of Stonnington, and the City of Whitehorse (Figure S1). This area collectively houses about 1.27 million inhabitants (Australian Bureau of Statistics, https://dbr.abs.gov.au) and covers approximately 650 km2, with slightly over 10% of that area being open greenspace.

The CNC evaluation for this node revolved around three pillars: (1) assessment of the local knowledge gained from the biodiversity records contributed during the CNC; (2) analysis of the CNC participants’ engagement; and (3) evaluation of the use of available greenspace by CNC participants.

To assess whether the CNC resulted in significant local knowledge gain, we summarised the CNC findings in light of existing biodiversity knowledge (Figure 3), represented by biodiversity records previously available in open-source, global biodiversity repositories for the same area.

**Figure 3.**
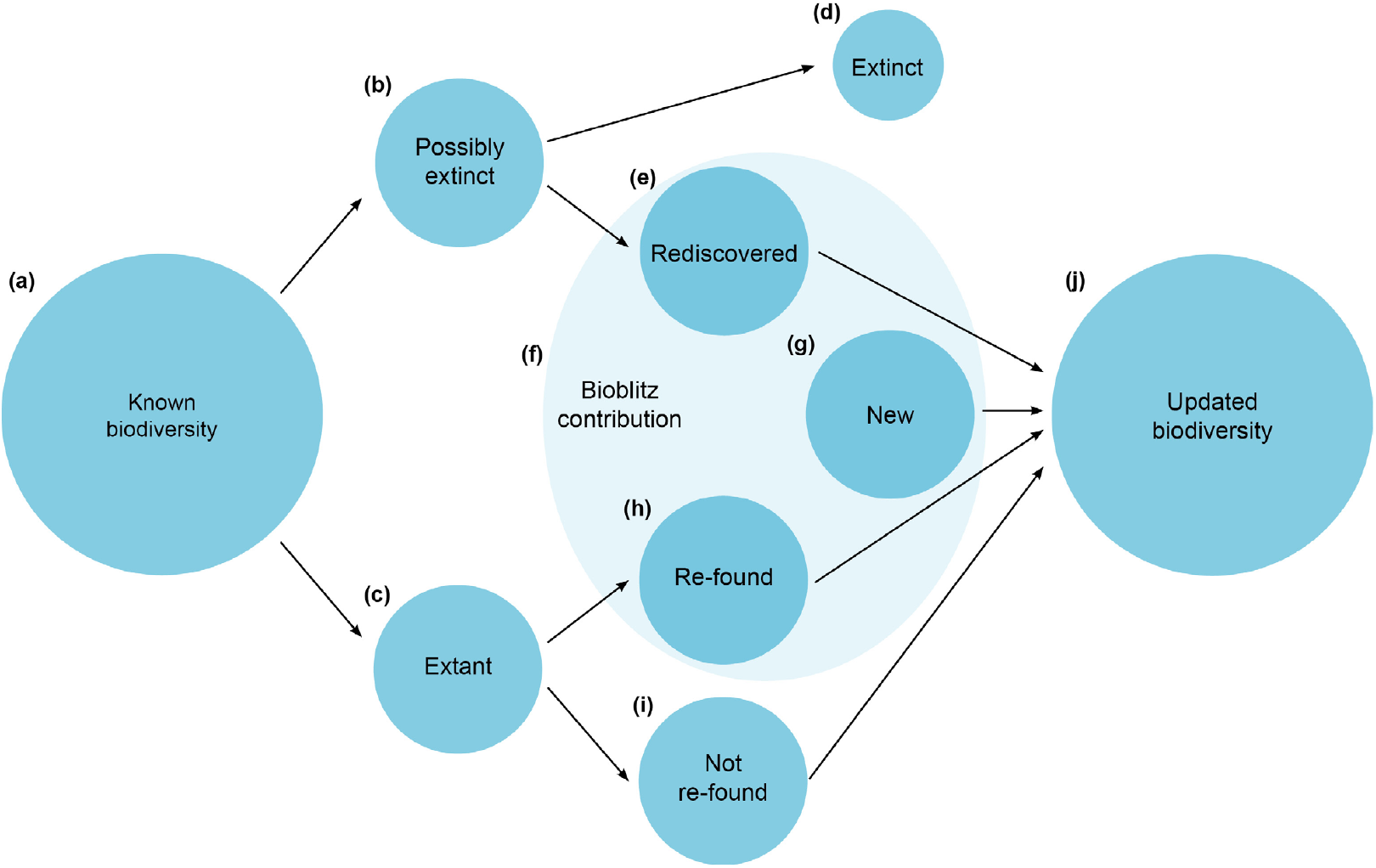
Conceptual framework of the contribution of citizen science events to local biodiversity knowledge. **(a)** Biodiversity known to the area before citizen science activity (species with historical and/or recent records on biodiversity repositories). **(b)** Possibly extinct species (species with historical but no recent records on biodiversity repositories). **(c)** Extant species (species with recent records on biodiversity repositories, with or without historical records). **(d)** Extinct species (species for which records will not be found anymore because they have gone locally extinct). **(e)** Rediscovered species (species with historical, but no recent, records - thought to be possibly extinct - for which records have been found during citizen science activity). **(f)** Species found during citizen science activity. **(g)** Newly discovered species (species without neither historical nor recent records that were found during citizen science activity). **(h)** Re-found species (extant species, with recent records on biodiversity repositories, found during citizen science activity). **(i)** Not re-found species (extant species, with recent records on biodiversity repositories, missed during citizen science activity). **(j)** Updated biodiversity known to the area after citizen science activity.

To analyse CNC participants’ engagement, we first looked at the contributions made by different participant types (i.e. members of the public vs. CNC organisers). Then, we checked the degree to which the already existing iNaturalist community contributed records to the CNC, and the growth of the iNaturalist community during the CNC.

Finally, we evaluated the use of greenspace across the ‘Melbourne Eastern Metropolitan Area’ node by CNC participants. To do this, we estimated the percentage of green-spaces where at least one record was collected, and then investigated whether the probability of greenspaces being visited depended on greenspace size.

A detailed explanation of our methodological approach, including data sourcing, statistical models and code is given in the Supplementary Material (Appendix B).

Even though biodiversity repositories point to an overwhelming interest in documenting birds over other taxonomic groups, CNC participants recorded a high diversity for other groups, in particular plants, insects, arachnids and fungi (Table S4). The records from the ‘Melbourne Eastern Metropolitan Area’ node also showed that the highest number of rediscovered and newly recorded species belong to these taxonomic groups. These findings indicate that the CNC provides a way to gather records for taxonomic groups that traditionally have poor representation across global biodiversity repositories, especially in terms of recent records (e.g. insects; Figure 4). Although the differences were subtle, the analytical framework proposed in this work (Figure 3) provides an updated picture of local biodiversity with a better representation of traditionally understudied groups (e.g. arachnids, fungi; Figure 4) and a slightly larger proportion of species introduced to Victoria (Figure S2).

**Figure 4.**
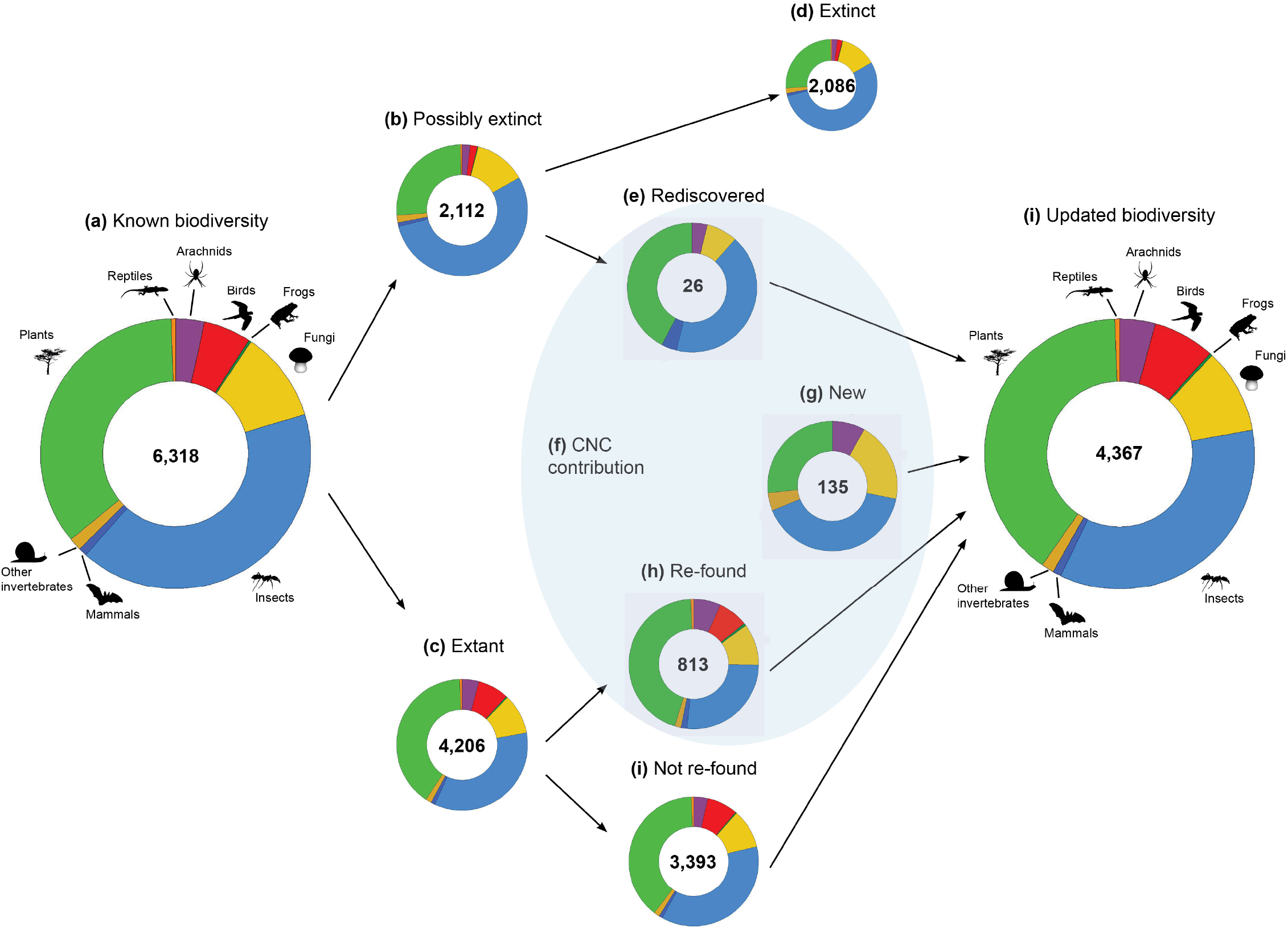
Contribution of the observations collected during the 2021 City Nature Challenge to local biodiversity knowledge from the Eastern Metropolitan Melbourne site node. See Table S4 for summaries of each taxonomic group.

Using the biodiversity records collected by the participants in the ‘Melbourne Eastern Metropolitan Area’ node, we also investigated different aspects of participant engagement. Out of the 291 citizens who collected records across this node, about 10% were organisers – including council officers, and experts co-running the events – and 90% were members of the public. Participants showed large variation in the number of records and species they reported, with CNC organisers contributing around 10 times more than members of the public (Table S5, Figure 5a-b).

**Figure 5.**
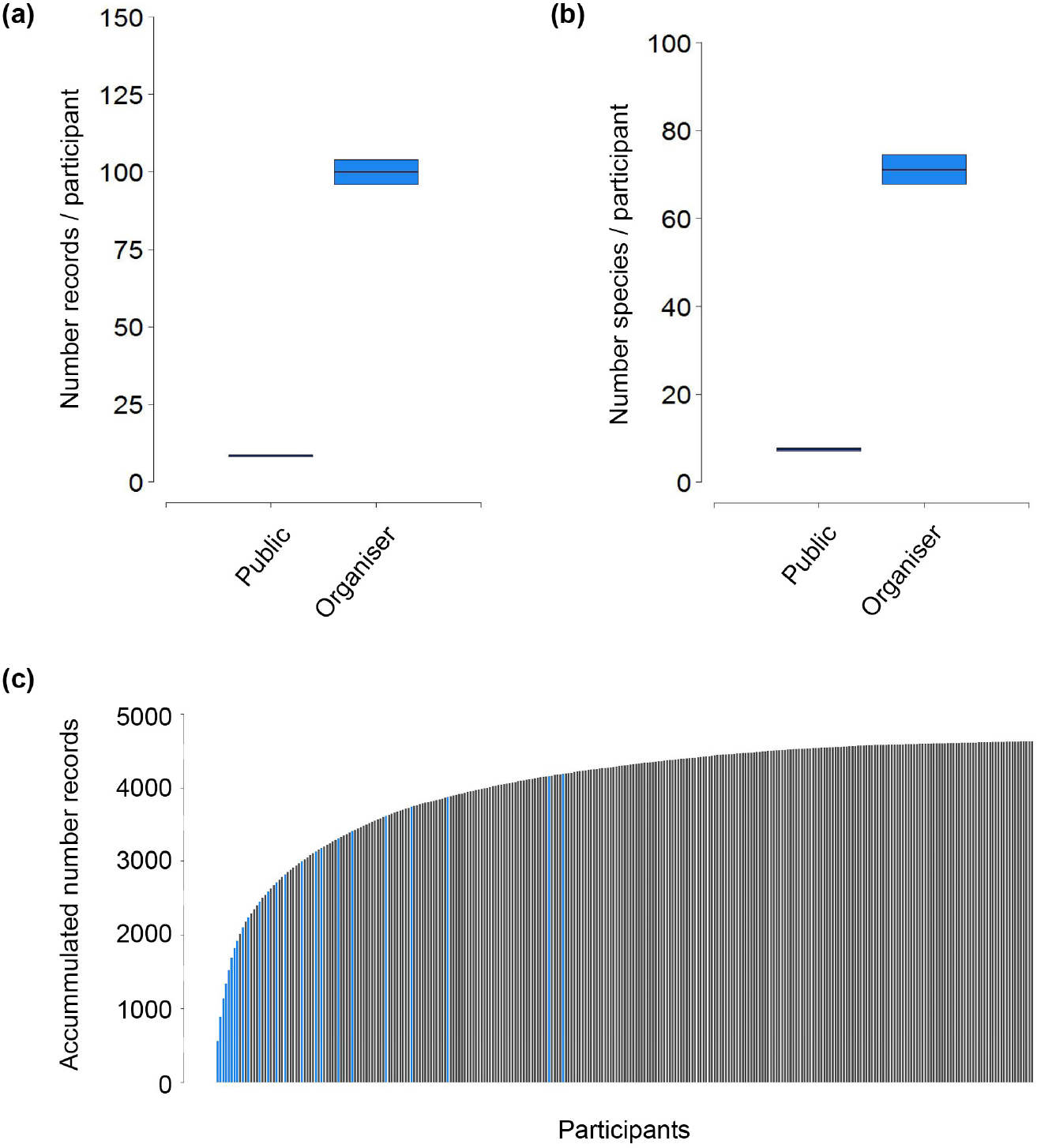
Participants’ contribution to the 2021 City nature Challenge across the Eastern Metropolitan Melbourne Area node. **(a)** Number of records contributed by different types of participants (members of the general public vs. CNC organisers). **(b)** Number of species contributed by different types of participants (members of the general public vs. CNC organisers). **(c)** Accumulated number of records contributed by participants. Participants are ordered from largest to smallest contribution in number of records, with members of the general public (n=267) shown in black and CNC organisers (n=24) shown in blue.

While over half of the participants contributed five or less records, eight participants only - all of them CNC organisers - contributed over a third of the totality of records (Figure 5c). Consequently, the contribution of records by participants was overall skewed towards small values.

Then, we turned our attention to the participants of the ‘Melbourne Eastern Metropolitan Area’ node and their relationship with the iNaturalist data collection platform (Table S6). Around 5% of existing iNaturalist users (i.e. those who had previously contributed records to this area) also contributed records during the CNC. Furthermore, the number of local iNaturalist users increased almost 8% during the same period (Table S5).

Finally, we focused on the biodiversity records that participants of the ‘Melbourne Eastern Metropolitan Area’ node collected from urban greenspaces and examined whether their collection was spatially uniform. We found that participants did not contribute observations evenly across greenspaces, both within and across councils (Table S5, Figure 6a-b). Participants collected one or more records from slightly over a quarter of the greenspaces across the node (Table S5, Figure 6a). Overall, the probability that participants visited individual greenspaces increased heavily with the size of the greenspace (Table S5), with more records being contributed from larger greenspaces – while the probability that participants visited greenspaces smaller than 100 m2 was close to zero, at least a fifth of greenspaces above 1,000,000 m2 were visited by the participants (Figure 6c).

**Figure 6.**
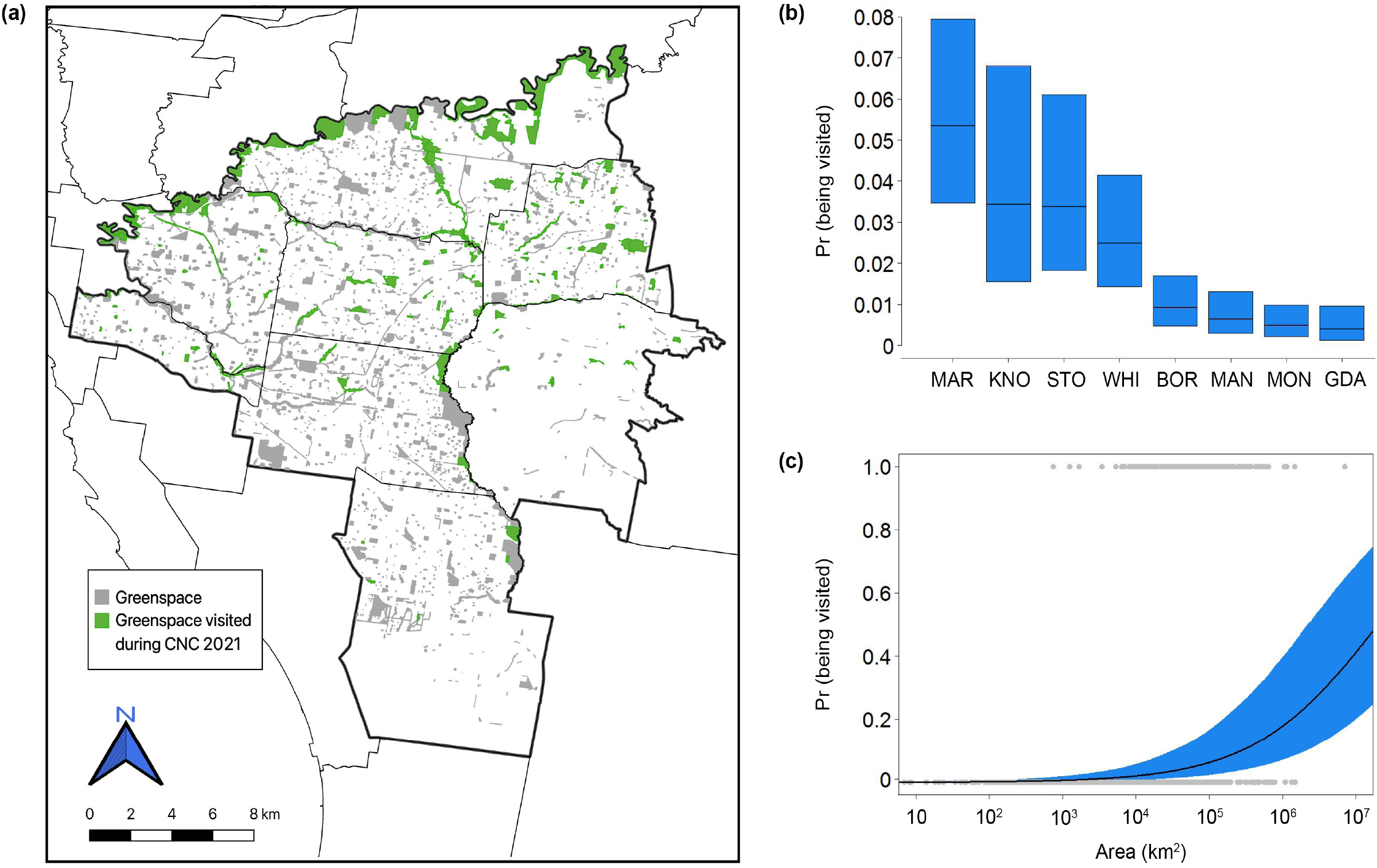
**(a)** Greenspace visited during the 2021 City Nature Challenge (in green) from all the available greenspace (in grey) across the Eastern Metropolitan Melbourne Area node. **(b)** Probability of greenspace to be visited across the eight city councils that formed the Eastern Metropolitan Melbourne Area node during the 2021 City Nature Challenge. **(c)** Probability of greenspace to be visited depending on the greenspace size.

The unstructured and opportunistic nature of biodiversity records collected during the CNC likely resulted in some spatial and taxonomic biases within the records contributed by the participants of this node (Geldmann et al. 2016, Mesaglio and Callaghan 2021). Notably, more than half of the observations were recorded from a handful of large greenspaces. This trend was likely exacerbated by the fact that the biodiversity surveys organised by the ‘Melbourne Eastern Metropolitan Area’ node were largely run from well-known, large greenspaces. What is more, the expertise of the organisers and/ or facilitators of these activities likely resulted in higher contribution of records and species for particular taxonomic groups they were experts on. For example, the involvement of the Entomological Society of Victoria in several events certainly drove the high contribution of insect records (Figure 4e,g-h). Reporting preferences toward common or more ‘familiar’ species, as well as those easier to detect, have been previously highlighted in the citizen science literature (Di Cecco et al. 2021, Johnston et al. 2022). In the ‘Melbourne Eastern Metropolitan Area’ node, 20 of the most highly reported 25 individual species were birds or plants. Additionally, the timing of the CNC may have an impact on the diversity of the taxa found for some groups, especially in the southern hemisphere – while fungi abound in April, other groups like grasses and herbs may not be easily found or identified at this time of the year. Running other citizen science activities complementary to the CNC can help to overcome this limitation. Particularly, the Great Southern Bioblitz (https://www.greatsouthernbioblitz.org/), which runs towards the end of the year and shares methodology and platforms with the CNC, represents an excellent opportunity to gain some synergy and amplify the diversity of taxa that citizen scientists can find in this node.

Beyond their contribution to understanding of local – and ultimately global –biodiversity, ongoing CNC-type events present local governments with cost-effective tools to make informed, evidence-based management and policy decisions. When citizen science initiatives are diverse (e.g. cover different areas within a management unit) and sustained through time, findings can support medium- and long-term conservation actions (Kobori et al. 2016). Examples include prompt protection of endangered or charismatic species (e.g. Park Victoria’s Data Discovery Program in Victoria, Australia), creation and monitoring of healthier habitats for humans and other species (e.g. Rakali as an indicator of river restoration success in Australia), planning for future climate scenarios, and early management of introduced species (e.g. European firebug in Melbourne, Australia; Mata et al. (2022)). Local governments and conservation groups are key actors in transforming citizen science findings into real, tangible management actions.

### Lessons learnt: Uptake of City Nature Challenge findings by local government

The ‘Melbourne Eastern Metropolitan Area’ node during the 2021 CNC was the first opportunity that many of the participating municipalities had in terms of organising and running an urban bioblitz event, and some of the aspirations and lessons learnt through this experience are shared here.

#### Education and engagement programs

One of the overriding benefits of the CNC bioblitz experience was the opportunity to facilitate activities that invited local people and communities to connect with nature through organised events, or by encouraging people to capture records as part of their day-to-day routine. Participants included a wide cross-section of the broader community, with young people, older people, and people from Culturally and Linguistically Diverse (CALD) backgrounds taking part; and organisers received positive feedback about their experiences working alongside biodiversity experts, and contributing in hands-on learning approaches. The use of the iNaturalist platform also opened up opportunities to connect with nature in a non-physical space, with features such as commenting and ‘agreeing’ with observations enabling access to biodiversity for other community members who had reduced capacity to visit the bushland environments in person for various reasons.

Another positive benefit was the increased exposure for local conservation groups and bushland reserves. Through the CNC bioblitz activities, the local community had a chance to become more aware of their local reserves and biodiversity, and for some people, particularly the CALD members of the community, the time spent in these places fostered a greater familiarity and appreciation for these sites. The experiences and greater familiarity is also likely to create deeper appreciation and passion for the local environments, which could translate to a more active network of citizens who may go on to participate in a wider range of programs, including wildlife gardening, nature strip planting, friends-of-groups, and other related programs focussing on sustainability and environmental stewardship.

The overriding hope is that events such as the CNC will help empower and enable residents to become stewards of biodiversity and seek out practical ways to continue to support local biodiversity in their gardens, streets, and local parks and reserves.

#### Citizen scientists

Participating in the CNC bioblitz provided individuals with the tools and knowledge to become more active citizen scientists and contribute high-quality data about the biodiversity intheir neighbourhood. Over time, there is the potential to create an expanded network of citizen scientists with a collective identity similar to other volunteering groups. These groups could self-organise or work in partnership with local councils to begin documenting biodiversity more strategically across time (e.g. seasons, or throughout the day or night) and space (e.g. surveying areas where there are information gaps), or to target particular species or groups. This will build a more comprehensive dataset that is updated more frequently than would be possible if local councils were commissioning surveys. Given that there are spikes in iNaturalist users during the CNC bioblitz events, and that the drop after the event remains at a higher baseline than prior to the event, there is a strong indication that this will become a reality for many local councils.

#### Municipality led conservation programs and applied research

The potential for bioblitz events to become a key source of information for local governments is still in the foundational stages, as events to date have largely been framed as community engagement and education activities that encourage connection to nature and provide an entry point into citizen science. However, as the citizen scientist movement grows there is increasing potential for them to contribute timely, targeted and high-quality records that can be used to inform policies and practices around management, education and other areas of co-developed research. This model of citizen science can also be used to monitor outcomes from projects to feed into adaptive management programs and evaluate success. The data collected can also be used as evidence to support funding requests, inform advocacy around key issues, and generally develop a deeper understanding of biodiversity in the local context.

The applied research and direct input of data into decision making is not just limited to biodiversity resilience plans; it can also be used for (1) planning, designing, improving and protecting urban greenspaces and habitat connectivity corridors; (2) informing climate response plans; and (3) monitoring and managing invasive species or biosecurity risks. Since multiple councils have collaborated during the CNC, there is also the opportunity for thinking and working at larger scales, pooling resources, and sharing learnings to accelerate progress towards improved outcomes.

#### Next steps towards future vision

While citizen scientist participation in urban bioblitzes such as the CNC have enormous potential to benefit biodiversity outcomes, there are still many gaps in our knowledge about how to maximise this potential. Some areas for additional investigation are presented below.

1. How do different approaches to event organisation influence the outcomes in terms of biodiversity records across time, space and taxa? This question can be investigated for local events, but really benefits from taking a larger picture look at events at the global scale. For example, what role does the participation of experts or friends-of-groups have on the type of data collected and long-term citizen science uptake or biodiversity stewardship? Can strategic messaging (e.g. a focus on reptiles or nature strips) or selection of event locations (e.g. parks or suburbs with very few records) help fill data gaps in a constructive way?
2. How can the bioblitz model be adjusted to allow for biodiversity records to be collected across different seasons? For example, the timing of the CNC is towards the end of the active period for biodiversity in the southern hemisphere, so participation in the Great Southern Bioblitz can help complement records collected during the CNC. Encouraging the continued use of iNaturalist to record incidental biodiversity sightings outside of the formal bioblitz, and empowering local groups to take on their own coordination of local collection events under the model described in the previous section will also help ensure records are collected over a broader spatial and temporal scale.
3. How long does it take to develop a consistent picture of biodiversity for an area through the bioblitz model of citizen science? Are there ways to accelerate this journey through activities identified in the previous steps, or by providing clear demonstrations of how data are currently being used to inform management practices or policies? This is where the real strength of the framework we present in Figure 3, and the analysis we present in Figures 4-6 comes to the fore.

### Scaling up to regional and global scales

We have shown that the contribution of biodiversity-based citizen science activities to the current biodiversity knowledge can be analysed through a simple, reproducible approach (Figure 3). Tools to evaluate the success of different dimensions of citizen science are urgently needed (Jordan et al. 2012, Kieslinger et al. 2018, Schaefer et al. 2021). Here, we demonstrate that the biodiversity data collected through bioblitz-type activities can be used to evaluate both their contribution to biodiversity knowledge and some aspects of participants’ engagement. Social metrics to evaluate satisfaction, learning or connectedness to nature, could complement this approach to provide a more comprehensive picture. The approach presented in this work can be readily taken up by, potentially, every participating node of the CNC on a given year, and across years, to evaluate nodes’ relative success. Moreover, the evaluation could be easily scaled up, with the totality of records collected yearly through the CNC compared against the totality of known biodiversity worldwide – e.g. as compiled in the Global Biodiversity Information Facility (https://www.gbif.org/).

Our shared experience of the 2021 CNC as a collaboration between city practitioners and biodiversity researchers has delivered enormous benefits for identifying a shared vision of how to maximise benefits from future bioblitz events. By working together and blending our individual knowledge and experiences we have begun to form a grander shared vision of the role that bioblitz style events can play in growing biodiversity knowledge, decision making and stewardship in the urban context. We look forward to working together on the next steps and sharing our progress with the broader global community.

## Acknowledgements

We would like to acknowledge the Traditional Owners of the land on which the ‘Melbourne Eastern Metropolitan Area’ node of the City Nature Challenge 2021 was organised, the Wurundjeri and Bunurong people of the Kulin Nation, and pay our respects to their Elders past, present and emerging, and honour their deep spiritual, cultural, and customary connections to the land. We extend our deepest appreciation to the Entomological Society of Victoria, the Field Naturalist Club of Victoria and all the Local Government Areas’ staff involved in organising and running the CNC events. We are thankful for all the CNC participants – this work would not have been possible without their enthusiasm. We thank the Natural History Museum of Los Angeles and the California Academy of Sciences as the original founders and organisers of the CNC and recognise the important role that iNaturalist and its funders have played in driving the CNC initiative to its global success.

## Data availability statement

Data and codes to reproduce models and plots are already published and publicly available in Zenodo: https://doi.org/10.5281/zenodo.7312210

## Notes

https://doi.org/10.5281/zenodo.7312210

### Competing Interest Statement

The authors have declared no competing interest.

