## Supplementary Material for "The City Nature Challenge – A global citizen science phenomenon contributing to biodiversity knowledge and informing local government practices"

**Appendix A:** iNaturalist

**Appendix B:** Analytical approach for the evaluation of the 'Melbourne Eastern Metropolitan Area' node of the 2021 City Nature Challenge

**Table S1.** Number of cities and countries taking part of the City Nature Challenge from 2016 to 2021, along with the approximated number of observations, species recorded and participants worldwide by year.

**Table S2.** Number of observations and participants in the 2021 City Nature Challenge by country.

**Table S3.** Summaries of biodiversity records for each of the eight city councils in the 'Melbourne Eastern Metropolitan Area' node, including historical observations, recent observations and observations made during the 2021 City Nature Challenge.

**Table S4.** Summaries of biodiversity records for each main taxonomic group found in the 'Melbourne Eastern Metropolitan Area' node, including historical observations, recent observations and observations made during the 2021 City Nature Challenge.

**Table S5.** Results from the 'Melbourne Eastern Metropolitan Area' node case study.

**Table S6.** Summary of iNaturalist use prior to and during the 2021 City Nature Challenge.

**Figure S1.** Location of the 'Eastern Metropolitan Melbourne Area' node of the 2021 City Nature Challenge.

**Figure S2.** Contribution of the observations collected during the 2021 City Nature Challenge to local biodiversity knowledge from the 'Eastern Metropolitan Melbourne Area' node by species' origin (i.e. indigenous or introduced to Victoria, Australia).

**Appendix A: iNaturalist**

iNaturalist (https://www.inaturalist.org/) is an online social platform designed map and share biodiversity observations across the world. Observations are uploaded as media, including photograph and audio files. Observations are publicly available but may only be contributed and curated by registered members. iNaturalist members uploading observations can choose between providing an taxonomical identifications themselves or leaving it to other iNaturalist community members to do so. iNaturalist is therefore both a crowdsourced tool to record biodiversity and curate species identifications, where collaboration amongst users is key to achieve the platform's maximum potential.

iNaturalist offers a voluntary, organic taxonomical curation of the data contributed to the platform. There is no limit to how many users can contribute identifications for a given record. For each observation, users can agree with previous identifications, help refine previous identifications to move them along until they reach species level, or correct previous mis-identifications. In general, the ability to identify contributed records will depend on the quality of uploaded media files. For some groups, media-based identifications will be limited to higher taxonomic levels, as lower taxonomical identification – for example to genus or species level – would require the direct observation and/or handling of collected specimens. Once identification consensus has been reached amongst, at least, two thirds of the users contributing identifications, the observation moves from 'Needs ID' to 'Research Grade', at which stage it flows programmatically to global repositories of biodiversity information such as the Global Biodiversity Information Facility (www. gbif.org). iNaturalist is also programatically linked with national-level repositories (e.g. Atlas of Living Australia; www.ala.org.au). Data collected through iNaturalist can be searched and downloaded using tools in these national and global repositories. Observations may also be classified as 'Casual', when they are (1) missing media and/or spatio-temporal metadata (e.g. date, locality) or (2) considered to be from a species that is either cultivated or in captivity.

**Appendix B: Analytical approach for the evaluation of the 'Melbourne Eastern Metropolitan Area' node of the 2021 City Nature Challenge**

*Data collection*

The City Nature Challenge 2021 (CNC for short) global event was held between April 30 and May 3, 2021. Biodiversity records collected within the 'Melbourne Eastern Metropolitan Area' node were uploaded to the iNaturalist project 'City Nature Challenge 2021: Melbourne Eastern Metropolitan Area', where they were curated by the iNaturalist community. We downloaded records of animals, plants and fungi a week after the CNC had finished (9 May 2021) and removed records classified as 'Casual'.

We also downloaded all animals, plants and fungi observations made before the CNC across the same study area and available from the Atlas of Living Australia (ALA for short). We categorised these records as ‘recent’ (made during the last 30 years – since 1 January 1991) or ‘historical’ (made longer than 30 years ago – before 1 January 1991).

We developed a programmatic pipeline to curate the CNC records along with the historical and recent records from ALA. We standardised the taxonomy following that used in ALA, and assigned main taxonomic group and origin (indigenous or introduced to Victoria) to each taxa. In cases where records lacked species-level identification, we assigned them to genus or family, and kept them as long as their genus or family was not represented amongst the species already identified in the full dataset.

*Statistical analyses*

The CNC evaluation comprised three complementary areas: (1) understanding the contribution of the CNC to local biodiversity knowledge; (2) engagement of participants; and (3) patterns of greenspace use.

1. Contribution to local biodiversity knowledge

We analysed the contribution of the CNC to local biodiversity knowledge using the combination of historical, recent and CNC records. For the whole ‘Melbourne Eastern Metropolitan Area’ node – as well as for each (1) individual municipality in the node; (2) main taxonomic group; and (3), species’ origin – we calculated the (i) number of historical, recent and CNC records and species; (ii) number of species with historical but no recent records; (iii) number of recent species that were or were not found during the CNC; (iv) number of species with historical but no recent records that were rediscovered during the CNC; and (v) number of species discovered for the first time during the CNC.

Next, using data at the council level, we ran linear models to estimate the (1) percentage of recent species that were re-found; (2) percentage of species that were re-discovered; (3) percentage of species that were newly discovered. These linear models were specified as:

$\text{Percentage}_i \sim \text{Normal}(\mu,\tau)$

where *i* represents a city council. The mean percentage estimate μ was given a Normal (0, 0.0001) prior and $\tau = 1/\sigma^2$, with σ having a Uniform (0, 100) prior. The percentage response variables were logit-transformed and standardised.

2. Participant engagement

Using the records collected during the CNC, we estimated the number of records and species contributed by participants, including different types of participants (CNC organisers vs. member of the public). We used regression models specified as:

$Count_i \sim Poisson(\lambda_i)$

$\lambda_i = exp(\alpha\_Participant\_type_i)$

where λ represents the estimated mean count by participant *i*, and the estimated mean count by different participant types (α_Participant_type) were given Normal (0, 0.01) priors.

In addition, using data at the council level, we ran linear models to estimate the (1) percentage of existing iNaturalist users (local to each city council) who contributed records during the 2021 CNC; and (2) percentage of iNaturalist users who contributed records for the first time during the 2021 CNC. We specified these linear models as:

$Percentage_i \sim Normal(\mu,\tau)$

where *i* represents a council, the mean percentage estimate μ was given a Normal (0, 0.0001) prior and $\tau = 1/\sigma^2$, with σ having a Uniform (0, 100) prior. The percentage response variables were logit-transformed and standardised.

3. Use of open greenspace

Using data at the council level, we estimated the percentage of available greenspace that was visited by CNC participants. This linear model was specified as:

$Percentage_i \sim Normal(\mu,\tau)$

where the response Percentage was calculated as the accumulated area of open greenspaces in council *i* where one or more records were collected during the 2021 CNC compared to the total area of open greenspaces in the same council. This percentage response variable was logit-transformed and standardised. The mean percentage estimate μ was given a Normal (0, 0.0001) prior and $\tau = 1/\sigma^2$, with σ having a Uniform (0, 100) prior.

We also estimated the probability that greenspaces were visited during the 2021 CNC, and whether greenspace size had an effect on that probability. We used an occupancy model of the form:

$Y_i \sim Bernoulli(p_i)$

$logit(p_i) = \alpha[Council_i] + \beta * Area_i$

$\alpha_j \sim$ Normal $(\mu,\tau)$

where the response variable Y represents whether greenspace *i* was or was not visited during the CNC, and p is the mean probability that greenspace *i* is visited. The parameter α represents the overall probability that greenspaces are visited, which is estimated for each council *j*. These values follow a Normal distribution, for which mean μ was given a Normal (0, 0.0001) prior and $\tau = 1/\sigma^2$, with σ having a Uniform (0, 100) prior. The parameter β represents the effect that greenspace area has on the probability that greenspaces are visited and was given a Normal (0, 0.0001) prior.

We estimated all model parameters under Bayesian inference, using Markov Chain Monte Carlo simulations. We implemented models in JAGS (Plummer 2003) through the R package R2jags (Yu-Sung and Masanao 2020). We used three chains of 100,000 iterations, discarding the first 20,000 in each chain as burn-in. For all models, convergence levels of R-hat < 1.1 were achieved (Gelman and Hill 2007).

Data and R code used in this work is publicly available at https://doi.org/10.5281/zenodo.7312210.

**Table S1.** Number of cities and countries taking part of the City Nature Challenge from 2016 to 2021, along with the approximated number of observations, species recorded and participants worldwide by year.

| Year | Number of cities | Number of countries | Number of observations | Number of species | Number of participants |
|---|---|---|---|---|---|
| 2016 | 2 | 1 | 19800 | 2500 | 1000 |
| 2017 | 16 | 1 | 125000 | 8600 | 4000 |
| 2018 | 68 | 17 | 441000 | 18000 | 17000 |
| 2019 | 159 | 28 | 963000 | 31000 | 32000 |
| 2020 | 244 | 40 | 815000 | 32600 | 41000 |
| 2021 | 419 | 44 | 1270000 | 45300 | 52000 |

**Table S2.** Number of observations and participants in the 2021 City Nature Challenge by country.

| Country | Number of observations | Number of participants |
|---|---|---|
| United States | 567129 | 32866 |
| South Africa | 108469 | 1838 |
| Mexico | 77832 | 1801 |
| Taiwan | 51415 | 1403 |
| United Kingdom | 51297 | 1827 |
| Canada | 48665 | 2712 |
| China | 38750 | 1376 |
| Austria | 33427 | 641 |
| Colombia | 22619 | 912 |
| New Zealand | 22465 | 580 |
| Australia | 20566 | 849 |
| Czech Republic | 20564 | 912 |
| Spain | 18295 | 483 |
| Brazil | 14182 | 595 |
| Russia | 13539 | 212 |
| Argentina | 13502 | 438 |
| Malaysia | 12943 | 531 |
| Panama | 12626 | 574 |
| Germany | 11855 | 321 |
| Italy | 10239 | 352 |
| Ecuador | 9303 | 306 |
| India | 7458 | 93 |
| Peru | 6770 | 434 |
| Ukraine | 5504 | 46 |
| Bolivia | 5403 | 117 |
| Luxembourg | 5341 | 268 |
| Japan | 5054 | 147 |
| Nicaragua | 4680 | 173 |
| Philippines | 4504 | 160 |
| Honduras | 3656 | 105 |
| Costa Rica | 2655 | 104 |
| Botswana | 2217 | 17 |
| Denmark | 2181 | 164 |
| El Salvador | 1148 | 64 |
| Netherlands | 775 | 37 |
| Algeria | 641 | 5 |
| Greece | 602 | 55 |
| Portugal | 600 | 120 |
| Dominican Republic | 594 | 20 |
| Nigeria | 455 | 9 |
| Croatia | 279 | 14 |
| Thailand | 273 | 34 |
| Slovakia | 240 | 20 |
| United Arab Emirates | 36 | 9 |

**Table S3.** Summaries of biodiversity records for each of the eight city councils in the 'Melbourne Eastern Metropolitan Area' node, including historical observations, recent observations and observations made during the 2021 City Nature Challenge.

| **Council** | **Number of observations [total]** | **Number of species [total]** | **Number of observations [historical]** | **Number of species [historical]** | **Number of observations [recent]** | **Number of species [recent]** | **Number of observations [CNC]** | **Number of species [CNC]** | **Number of recent species found during CNC** | **Number of recent species not found during CNC** | **Number of extinct species** | **Number of rediscovered species** | **Number of newly found species during CNC** | **Number of newly found exotic species during CNC** |
|---|---|---|---|---|---|---|---|---|---|---|---|---|---|---|
| Monash | 262,017 | 1,761 | 10,504 | 1,059 | 251,113 | 1,181 | 400 | 193 | 132 | 1,049 | 519 | 4 | 57 | 14 |
| Knox | 184,515 | 3,047 | 20,966 | 1,901 | 162,865 | 1,951 | 684 | 321 | 240 | 1,711 | 1,015 | 5 | 76 | 6 |
| Manningham | 152,824 | 2,889 | 24,811 | 1,425 | 127,634 | 2,412 | 379 | 213 | 175 | 2,237 | 439 | 4 | 34 | 4 |
| Greater Dandenong | 100,055 | 1,805 | 5,238 | 1,017 | 94,672 | 1,292 | 145 | 91 | 67 | 1,225 | 489 | 0 | 24 | 3 |
| Whitehorse | 97,622 | 2,553 | 13,330 | 1,367 | 83,320 | 1,868 | 972 | 389 | 312 | 1,556 | 608 | 9 | 68 | 11 |
| Boroondara | 44,755 | 1,770 | 14,012 | 1,159 | 30,500 | 1,085 | 243 | 121 | 83 | 1,002 | 647 | 8 | 30 | 3 |
| Maroondah | 33,512 | 2,295 | 8,430 | 1,491 | 23,422 | 1,274 | 1,660 | 475 | 291 | 983 | 837 | 14 | 170 | 9 |
| Stonnington | 18,791 | 935 | 1,194 | 497 | 17,442 | 571 | 155 | 102 | 54 | 517 | 316 | 3 | 45 | 18 |
| **Total** | 894,091 | 6,453 | 98,485 | 4,142 | 790,968 | 4,206 | 4,638 | 974 | 813 | 3,393 | 2,086 | 26 | 135 | 22 |

**Table S4.** Summaries of biodiversity data for each main taxonomic group found in the Melbourne Eastern Metropolitan Area, including historical observations, recent observations and observations made during the 2021 City Nature Challenge.

| Main group | Number of observations [total] | Number of species [total] | Number of observations [historical] | Number of species [historical] | Number of observations [recent] | Number of species [recent] | Number of observations [CNC] | Number of species [CNC] | Number of recent species found during CNC | Number of recent species not found during CNC | Number of extinct species | Number of rediscovered species | Number of newly found species during CNC | Number of newly found exotic species during CNC |
|---|---|---|---|---|---|---|---|---|---|---|---|---|---|---|
| Birds | 786,410 | 359 | 56,721 | 307 | 728,989 | 322 | 700 | 61 | 61 | 261 | 37 | 0 | 0 | 0 |
| Plants | 74,289 | 2,270 | 32,731 | 1,639 | 39,482 | 1,688 | 2,076 | 409 | 362 | 1,326 | 535 | 11 | 36 | 22 |
| Insects | 16,932 | 2,664 | 5,192 | 1,604 | 10,744 | 1,464 | 996 | 281 | 215 | 1,249 | 1,134 | 11 | 55 | 0 |
| Fungi | 4,743 | 711 | 694 | 355 | 3,702 | 414 | 347 | 113 | 84 | 330 | 268 | 2 | 27 | 0 |
| Mammals | 4,106 | 64 | 1,839 | 55 | 2,185 | 43 | 82 | 13 | 12 | 31 | 20 | 1 | 0 | 0 |
| Frogs | 3,694 | 19 | 499 | 15 | 3,170 | 15 | 25 | 5 | 5 | 10 | 4 | 0 | 0 | 0 |
| Arachnids | 2,105 | 229 | 309 | 86 | 1,542 | 175 | 254 | 67 | 55 | 120 | 42 | 1 | 11 | 0 |
| Reptiles | 1,032 | 34 | 311 | 30 | 704 | 25 | 17 | 6 | 6 | 19 | 9 | 0 | 0 | 0 |
| Other invertebrates | 780 | 103 | 189 | 51 | 450 | 60 | 141 | 19 | 13 | 47 | 37 | 0 | 6 | 0 |
| **Total** | 894,091 | 6,453 | 98,485 | 4,142 | 790,968 | 4,206 | 4,638 | 974 | 813 | 3,393 | 2,086 | 26 | 135 | 22 |

**Table S5.** Model estimates for the 'Melbourne Eastern Metropolitan Area' node analyses.

| | Mean | 95% CI |
|---|---|---|
| **Contribution to local biodiversity knowledge** | | |
| Percentage of recent species found during CNC | 10.6% | 6.5% - 16.8% |
| Percentage of species newly discovered during CNC | 3.7% | 1.9% - 7.0% |
| Percentage of species re-discovered during CNC | 1.0% | 0.6% - 1.5% |
| **Participant engagement** | | |
| Number of records contributed by participant \| type = CNC organiser | 99.9 | 98.5 - 104.0 |
| Number of records contributed by participant \| type = general public | 8.4 | 8.3 - 8.7 |
| Number of species contributed by participant \| type = CNC organiser | 71.1 | 67.8 - 74.6 |
| Number of species contributed by participant \| type = general public | 7.4 | 7.0 - 7.7 |
| Percentage previous iNaturalist users that participated in CNC | 5.4% | 3.3% - 8.8% |
| Percentage increase in iNaturalist users during CNC | 7.7% | 4.3% - 13.7% |
| **Open greenspace use** | | |
| Percentage of total greenspace area visited during CNC | 29% | 15% - 47% |
| Probability of greenspace to be visited during CNC \| council = MAR | 0.053 | 0.035 - 0.079 |
| Probability of greenspace to be visited during CNC \| council = KNO | 0.034 | 0.016 - 0.068 |
| Probability of greenspace to be visited during CNC \| council = STO | 0.034 | 0.018 - 0.061 |
| Probability of greenspace to be visited during CNC \| council = WHI | 0.025 | 0.014 - 0.041 |
| Probability of greenspace to be visited during CNC \| council = BOR | 0.009 | 0.007 - 0.017 |
| Probability of greenspace to be visited during CNC \| council = MAN | 0.006 | 0.003 - 0.013 |
| Probability of greenspace to be visited during CNC \| council = MON | 0.005 | 0.002 - 0.010 |
| Probability of greenspace to be visited during CNC \| council =GDA | 0.004 | 0.001 - 0.010 |
| Probability of greenspace to be visited during CNC \| effect of greenspace size | 1.86 | 1.61 - 2.12 |

**Table S6.** Summary of iNaturalist use prior to and during the 2021 City Nature Challenge.

| Council | Number of iNaturalist users prior to CNC | Number of iNaturalist users who contributed during CNC | Number of new iNaturalist users during CNC | Percentage of former iNaturalist users who participated in CNC | Percentage increase in iNaturalist users during CNC |
|---|---|---|---|---|---|
| Boroondara | 436 | 41 | 29 | 2.75 | 6.24 |
| Greater Dandenong | 257 | 19 | 7 | 4.67 | 2.65 |
| Knox | 320 | 40 | 26 | 4.38 | 7.51 |
| Manningham | 417 | 43 | 28 | 3.60 | 6.29 |
| Maroondah | 197 | 80 | 53 | 13.71 | 21.20 |
| Monash | 411 | 42 | 22 | 4.87 | 5.08 |
| Stonnington | 244 | 37 | 24 | 5.33 | 8.96 |
| Whitehorse | 308 | 84 | 53 | 10.06 | 14.68 |

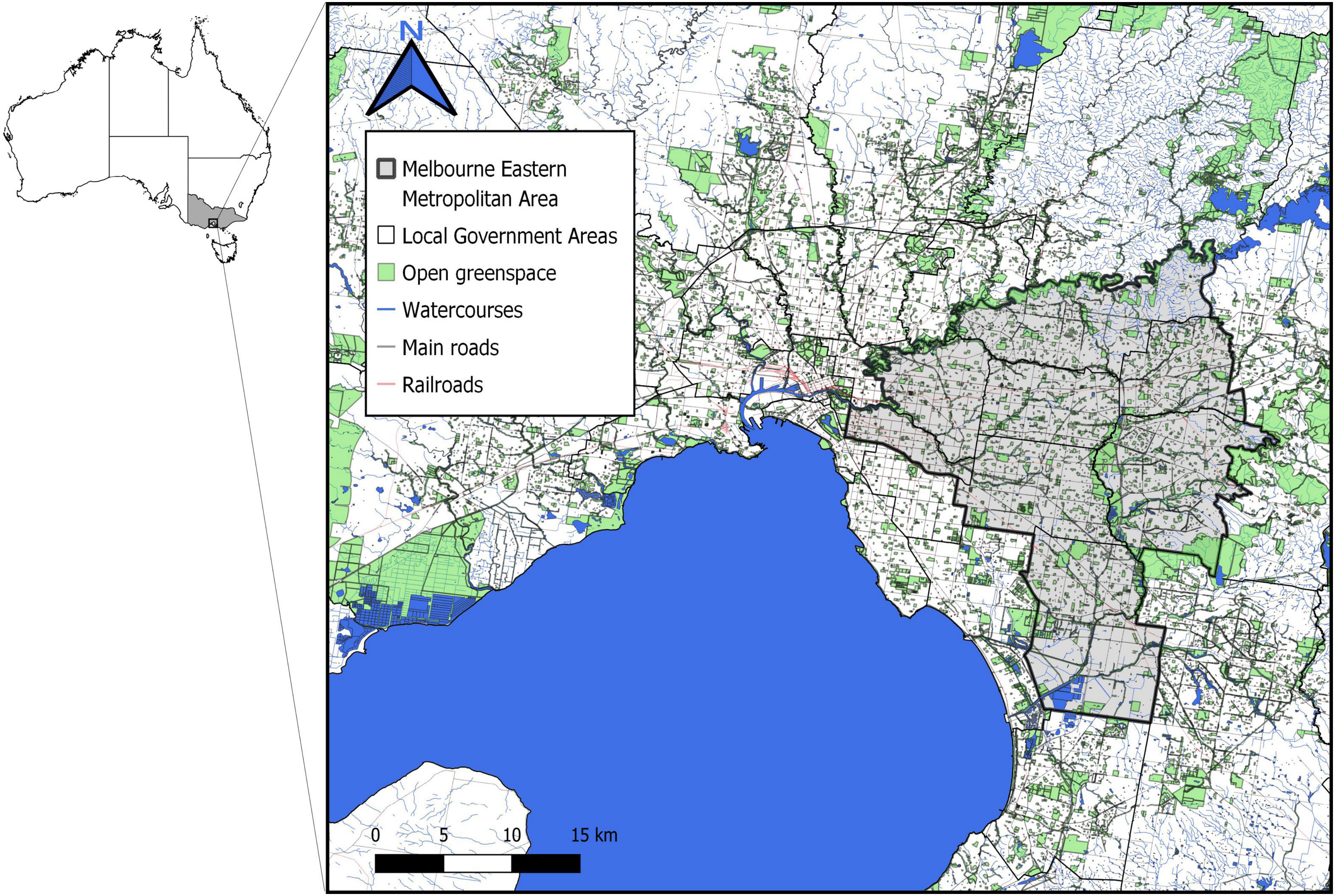


**Figure S1.** Location of the 'Eastern Metropolitan Melbourne Area' node of the 2021 City Nature Challenge.

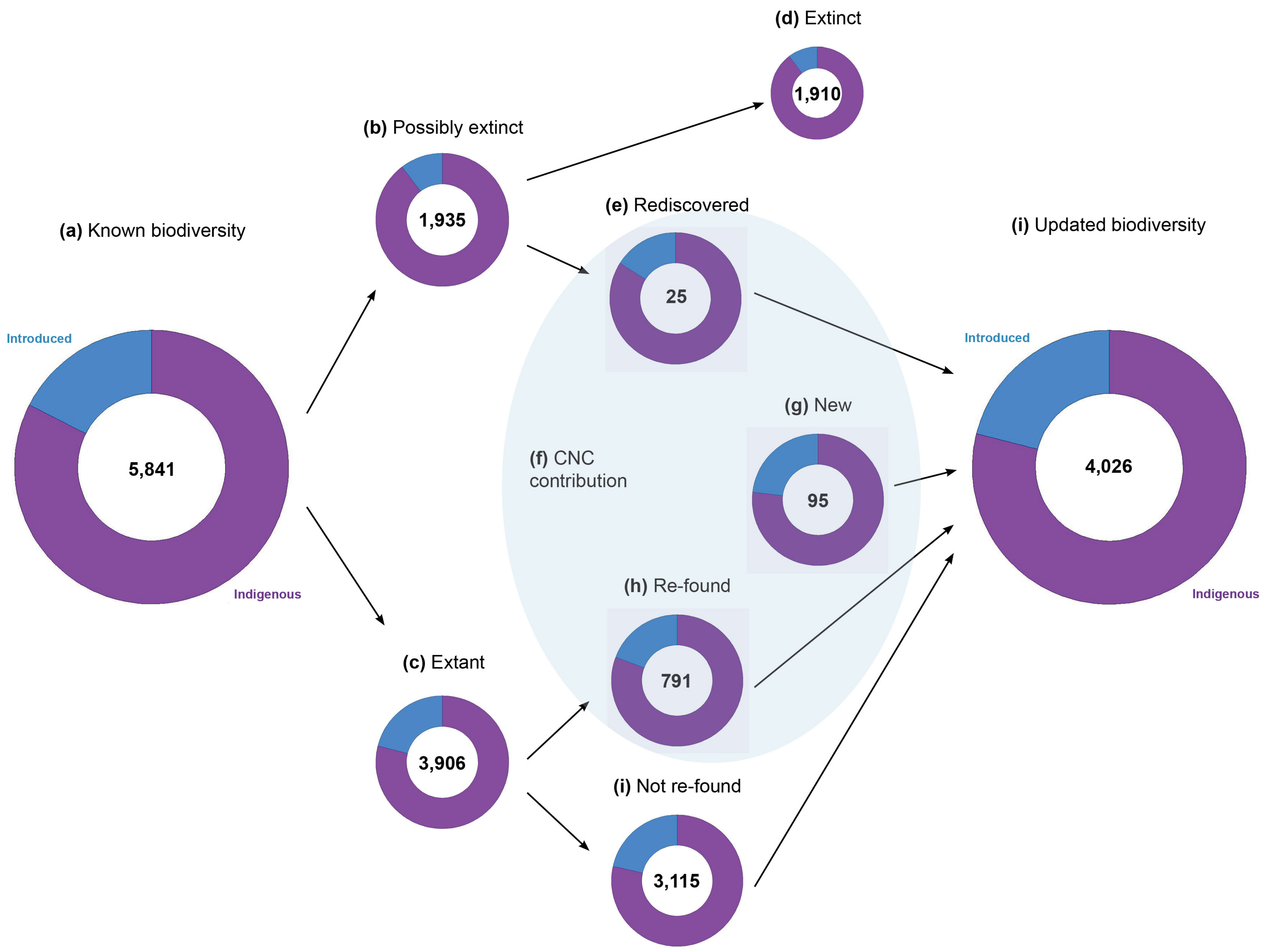


**Figure S2.** Contribution of the observations collected during the 2021 City Nature Challenge to local biodiversity knowledge from the 'Eastern Metropolitan Melbourne Area' node by species' origin (indigenous or introduced to Victoria, Australia).
